# Sampling strategies and pre-pandemic surveillance gaps for bat coronaviruses

**DOI:** 10.1101/2022.06.15.496296

**Authors:** Lily E. Cohen, Anna C. Fagre, Binqi Chen, Colin J. Carlson, Daniel J. Becker

**Affiliations:** Icahn School of Medicine at Mount Sinai, New York, NY, USA; Department of Microbiology, Immunology, and Pathology, College of Veterinary Medicine and Biomedical Sciences, Colorado State University, Fort Collins, CO, USA; Bat Health Foundation, Fort Collins, CO, USA; Center for Global Health Science and Security, Georgetown University Medical Center, Washington, D.C., USA; Department of Biology, University of Oklahoma, Norman, OK, USA

**Keywords:** Coronaviridae, bats, longitudinal sampling, viral ecology, open data

## Abstract

The emergence of SARS-CoV-2, and the challenge of pinpointing its ecological and evolutionary context, has highlighted the importance of evidence-based strategies for monitoring viral dynamics in bat reservoir hosts. Here, we compiled the results of 93,877 samples collected from bats across 111 studies between 1996 and 2018, and used these to develop an unprecedented open database, with over 2,400 estimates of coronavirus infection prevalence or seroprevalence at the finest methodological, spatiotemporal, and phylogenetic level of detail possible from public records. These data revealed a high degree of heterogeneity in viral prevalence, reflecting both real spatiotemporal variation in viral dynamics and the effect of variation in sampling design. Phylogenetically controlled meta-analysis revealed that the most significant determinant of successful viral detection was repeat sampling (i.e., returning to the same site multiple times); however, fewer than one in five studies longitudinally collected and reported data. Viral detection was also more successful in some seasons and from certain tissues, but was not improved by the use of euthanasia, indicating that viral detection may not be improved by terminal sampling. Finally, we found that prior to the pandemic, sampling effort was highly concentrated in ways that reflected concerns about zoonotic risk, leaving several broad geographic regions (e.g., South Asia, Latin America and the Caribbean, and most of Sub-Saharan Africa) and bat subfamilies (e.g., Stenodermatinae and Pteropodinae) measurably undersampled. These gaps constitute a notable vulnerability for global health security and will likely be a future barrier to contextualizing the origin of novel zoonotic coronaviruses.

## Introduction

Since the emergence of severe acute respiratory syndrome-associated coronavirus (SARS-CoV) in 2002, coronaviruses (Coronaviridae: Orthocoronavirinae) have been the subject of concern as potential pandemic threats. The group comprises four genera containing an estimated hundreds or thousands of viruses [1]. Two of these genera, the delta- and gammacoronaviruses, are primarily pathogens of birds, though they infect a handful of mammals: notably, porcine deltacoronavirus became the first shown to infect humans in 2021 [2]. The alpha- and betacoronaviruses contain all other known human-infective coronaviruses; the latter includes SARS-CoV, Middle East respiratory syndrome–related coronavirus (MERS-CoV), and severe acute respiratory syndrome coronavirus 2 (SARS-CoV-2), the three highly pathogenic coronaviruses that have caused significant morbidity and mortality in humans [3]. While alpha- and betacoronaviruses exhibit a high degree of host plasticity, there is substantial diversity of these viruses in bats, which are likely the ancestral hosts of these groups [4,5]. As such, coronaviruses have been among a handful of other clades of zoonotic pathogens (e.g., filoviruses, lyssaviruses, and henipaviruses) that have been monitored extensively in wild bats, and continue to be the subject of ongoing surveillance [6].

Research into the natural origins of SARS-CoV-2, and a broader renewed interest in coronavirus ecology and evolution, have highlighted the immense value of these surveillance studies. However, outside of long-term coordinated research projects, field sampling is often opportunistic in response to concerns about spillover, and capacity for systematic sampling is frequently financially- or logistically-constrained [7]. For example, prior comparative analyses of bat filovirus and henipavirus positivity have found that only a small fraction of studies report longitudinal data, limiting inference into temporal dynamics of infection in bats [6]. In turn, this limits the interpretability of these data in aggregate: for example, single sampling events can bias prevalence estimates in biologically meaningful ways (e.g., if sampling is more convenient in one season over another), and may lead to non-randomly missing data. In contrast, explicit spatiotemporal sampling designs can identify seasonal and environmental drivers of viral prevalence and shedding intensity, but these are logistically challenging and can necessitate prioritizing either spatial or temporal replication at the expense of the other scale [6]. These are essential considerations for study design, particularly if the ultimate goal is to explain and predict pathogen spillover, a dynamic process that is driven by geographical and temporal variation in infection prevalence and shedding from reservoir hosts [6,8], and the relative importance of non-spatiotemporal factors that may impact virus positivity (e.g., tissues sampled, use of euthanasia, diagnostic method) further warrants examination. Presently, our ability to quantify whether and how these factors shape global assessments of coronavirus spillover risk is limited by a lack of standardized and aggregated data from disparate studies.

Here, we compiled a standardized global database of infection prevalence and seroprevalence estimates from pre-pandemic coronavirus testing in wild bats, alongside relevant metadata on bat and viral taxonomy, study methodology, bat demography and seasonality, and ecological context. We first identified global biases in the distribution and intensity of pre-pandemic bat coronavirus surveillance, followed by comparative analyses to quantify phylogenetic signal in sampling effort and identify especially oversampled or undersampled bat clades. Next, we used a phylogenetically controlled meta-analysis to identify study designs, spatiotemporal factors, and biological traits that predict higher viral prevalence, with the aim of identifying potential ways to optimize future sampling. More broadly, we evaluate the global state of coronavirus surveillance in natural bat hosts prior to SARS-CoV-2-motivated research efforts.

## Results

### Descriptive analyses

From publicly available literature over the last quarter-century, we were able to recover data on 93,877 tests worth of coronavirus surveillance in bats. Over 90% of the 2,434 data points in our database report infection prevalence (93.7%; compared to 6.3% seroprevalence data ascertained using a mix of immunologic assays, including ELISA, western blot, and indirect immunofluorescence). Within the pooled-coronavirus genera (i.e., alpha- and betacoronavirus) infection prevalence dataset, nearly 95% of estimates used PCR targeting the RNA-dependent RNA polymerase (RdRp) gene; other gene targets included subunits of the coronavirus spike protein, the nucleocapsid gene, or the envelope protein. Of the 99.6% of rows detecting coronaviruses via PCR, approximately 56% used single-round PCR as opposed to nested PCR or multiple PCR assays in parallel (e.g., targeting different genes on the same RNA sample). More than half of these records (53.8%) based their primers on protocols from four past studies [9–12]. 34.8% of the pooled-coronavirus genera infection prevalence records were derived from studies that had euthanised their sampled bats. Table S2 shows the distribution of tissue types analyzed and the associated percentages of positive and zero infection prevalence values. Fecal samples and rectal swabs were the most common tissue used to detect coronavirus RNA. Sex and/or reproductive status of the bats sampled was only described in 12.6% of studies (14/111), resulting in 10% of individual prevalence records being stratified by sex.

### Spatial bias in surveillance effort

Prior to the COVID-19 pandemic, we found recoverable data describing sampling of wild bats for coronaviruses across 54 countries spanning six continents. However, we found that the distribution and intensity of viral surveillance has been starkly uneven (Fig. 1). Sampled countries varied in having one to 32 bat coronavirus studies (Fig. 1a), with the number of total samples tested ranging from four to 26,313 (Fig. 1b). Whereas sampling has occurred across all North American countries, both Central America and South America have had sparse surveillance. Similarly, sampling in sub-Saharan Africa as well as Central and South Asia has been inconsistent, with the majority of global surveillance having taken place in China, and to a lesser extent other regions of Southeast Asia. A generalized linear model (GLM) of binary sampling effort (*χ*^2^ = 12.08, *p* = 0.02, *R*^2^ = 0.04) confirmed that countries in Asia and Europe were marginally more likely to be sampled for bat coronaviruses than those in the Americas (Table S3). We found more substantial geographic biases regarding the relative intensity of sampling, specifically from the number of studies (*χ*^2^ = 17.08, *p* = 0.002, *R*^2^ = 0.05) and the number of tested samples (*χ*^2^ = 19549, *p* < 0.001, *R*^2^ = 0.11). Post-hoc comparisons from GLMs revealed significantly more studies per country in Asia compared to Africa and to Europe (Table S4). Similarly, the greatest contrast in total number of tested samples was between Asia and Europe (risk ratio [RR] =4.41) and between the Americas and Europe (RR = 2.11; Table S5).

**Figure 1.**
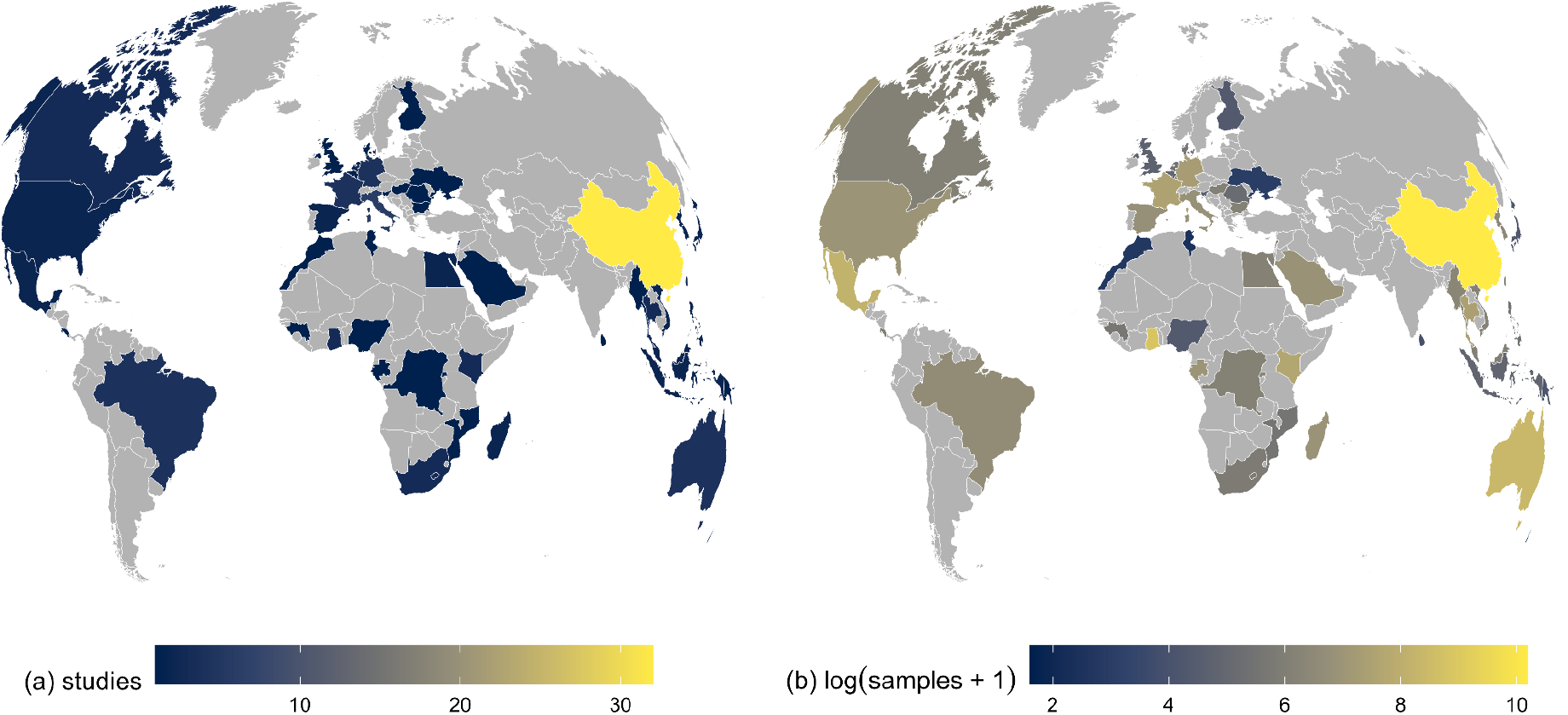
Geographic distribution of bat coronavirus sampling effort, defined by the number of studies per country (a) and the number of samples tested per country (b). Sampled countries varied in having one to 32 bat coronavirus studies (a), with the number of total samples tested ranging from four to 26,313 (b). A disproportionate number of bat coronavirus studies and testable samples were conducted and assayed in China, likely reflecting interest in the subgenus *Sarbecovirus* and the risk of future SARS-like virus emergence. Many areas were severely understudied, particularly relative to ecological and evolutionary risk factors for emergence [19]. In particular, sampling in Central and South America, sub-Saharan Africa, and Central and South Asia was notably limited.

### Taxonomic biases in surveillance effort

Over one in four bat species (363 species of the 1,287 included in our phylogeny [13]) were at some point targeted by pre-pandemic coronavirus surveillance. Surprisingly, bats have been sampled relatively evenly across the phylogeny (Fig. 2a). Indeed, we only identified intermediate phylogenetic signal in binary sampling effort (*D* = 0.88) that departed from both phylogenetic randomness (*p* < 0.001) and Brownian motion models of evolution (*p* <0.001). Similarly, phylogenetic factorization [14], a graph-partitioning algorithm based on the bat phylogeny, did not identify any bat clades that differed significantly in their fraction of sampled species. In contrast, we observed stronger taxonomic biases in sampling intensity. The number of studies per sampled species ranged from one to 24 (*Miniopterus schreibersii*), whereas the number of total samples tested ranged from one to 16,628 (*Rhinolophus sinicus*). The number of studies per sampled species showed low phylogenetic signal (λ = 0.04) that departed from Brownian motion models of evolution (*p* < 0.001) but not phylogenetic randomness (*p* = 0.35); phylogenetic factorization did, however, more flexibly identify four bat clades with significantly greater mean numbers of studies than the paraphyletic remainder (Fig. 2b): a subclade of the genus *Myotis* (including both European and Asian species), a subclade of the tribe Pipistrellini (including pipistrelle and noctule bats), the sister families Hipposideridae and Rhinolophidae, and the whole genus *Miniopterus* (Table S8).

**Figure 2.**
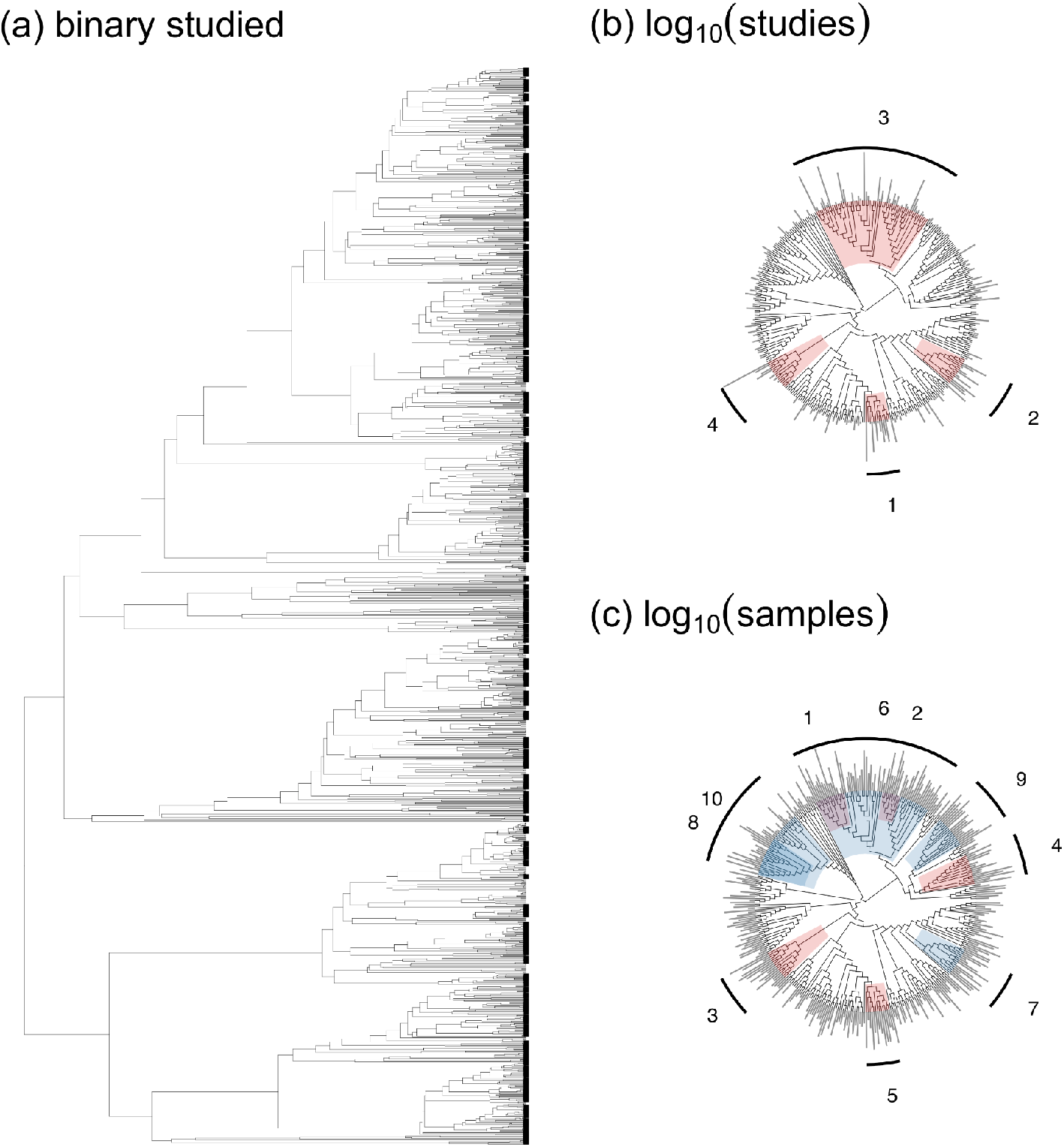
Evolutionary distribution of bat coronavirus sampling effort, defined as whether a bat species has been sampled (a), the number of studies (b), and the number of samples tested (c). Clades identified by phylogenetic factorization with greater or lesser sampling effort compared to a paraphyletic remainder are shown in red and blue, respectively, alongside clade numbers per analysis. Phylogenetic factorization did not identify any taxonomic patterns in binary sampling effort across the bat phylogeny (a) but did identify a number of bat clades within sampled bat species that have been particularly well-sampled for coronaviruses, both in terms of number of studies (b; Table S8) and number of samples (c; Table S9, only the first 10 phylogenetic factors are displayed). For analyses of total studies and tested samples, segment length corresponds to the relative degree of sampling effort.

For the total number of tested samples per species, we instead observed more intermediate phylogenetic signal (λ = 0.2) that departed from both Brownian motion models of evolution (*p* < 0.001) as well as phylogenetic randomness (*p* < 0.001). Accordingly, phylogenetic factorization identified a total of 23 clades with differential intensities of sampling effort, seven of which had relatively more tested samples and 16 of which had relatively fewer tested samples (Fig. 2c). The top clades with comparatively fewer total samples included the sister families Hipposideridae and Rhinolophidae as well as the above subclade of the tribe Pipistrellini, suggesting a greater number of publications on these bats but fewer tested samples. However, smaller subclades of the Hipposideridae and Rhinolophidae families were some of the most heavily sampled, suggesting key biases in sampling effort within these taxa that have been the subject of much coronavirus research (Table S9). Finally, members of the subfamily Stenodermatinae within phyllostomid bats were undersampled, as were several genera within the Pteropodinae subfamily (i.e., *Pteropus, Eidolon*, and *Acerodon*).

### Heterogeneity in coronavirus infection prevalence

Using a phylogenetic meta-analysis model that accounted for sampling variance, bat phylogeny, additional species effects, and within- and between-study variation [15,16], we observed high heterogeneity among coronavirus infection prevalence estimates (*I^2^* = 86.32%, *Q_2075_* = 12995.13, *p* < 0.0001). This heterogeneity was mainly driven by within-study (42.15%) and between-study effects (37%), with lesser contributions from bat phylogeny (7.04%) and additional species effects (0.13%). When repeating this intercept-only model for alphacoronavirus- and betacoronavirus-specific datasets, prevalence showed similar patterns of heterogeneity (alphacoronavirus: *I*^2^ = 82.37%, *Q_1769_* = 8759.34, *p* < 0.0001; betacoronavirus: *I^2^* = 76.9%, *Q_1626_* = 6043.81, *p* < 0.0001), driven primarily by within-study (alphacoronavirus: 46.53%; betacoronavirus: 36.43%) and between-study effects (alphacoronavirus: 29.003%; betacoronavirus: 27.10%), and secondarily by phylogenetic (alphacoronavirus: 6.83%; betacoronavirus: 13.37%) and other species-level effects (alphacoronavirus: 0.003%; betacoronavirus: 0.003%).

### Methodological and biological predictors of infection prevalence

When considering our suite of methodological and biological predictors in phylogenetic meta-analysis models, the fixed effects explained approximately 20% of the variance in infection prevalence (pooled-coronavirus genera *R^2^*: 0.21; alphacoronavirus-only *R^2^*: 0.21; betacoronavirus-only *R^2^*: 0.20). Across all three datasets, repeat sampling was associated with a 0.84-1.6% percentage point increase in coronavirus prevalence (pooled coronavirus: untransformed *β* = 0.15; 95% confidence interval (CI) 0.06-0.25, *p* < 0.005; alphacoronavirus: untransformed *β* = 0.14; 95% 0.03-0.26, p < 0.05; betacoronavirus: untransformed *β* = 0.14; 95% CI: 0.04-0.24, *p* < 0.05) as compared to one-time (single) sampling (Fig. 3). Similarly, longitudinal study design predicted a small increase (~ 0.2-0.3% percentage points) in positive viral detection in the pooled coronavirus (untransformed *β* = 0.06; 95% CI: 0.02-0.11, *p* < 0.01) and alphacoronavirus-only (untransformed *β* = 0.07; 95% CI: 0.02-0.12, *p* < 0.01) datasets, as opposed to cross-sectional sampling. Other model variables including tissue type, sampling season, bat family, PCR type, and gene target showed weak or no significant association with coronavirus positivity across all datasets. Notably, use of euthanasia was not associated with greater ability to detect coronavirus RNA.

**Figure 3.**
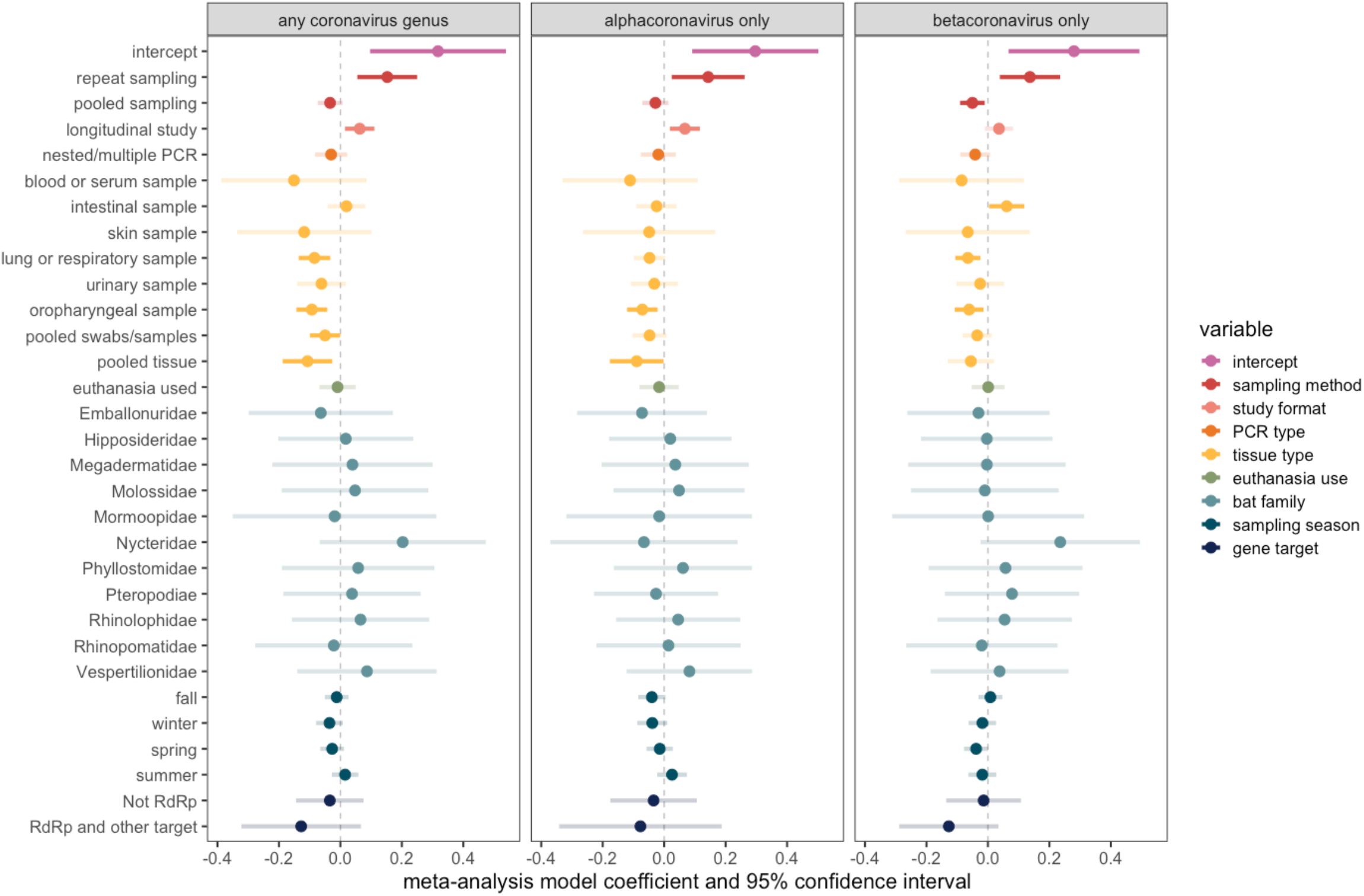
Methodological and biological predictors of coronavirus prevalence in wild bats. Phylogenetic meta-analysis model coefficients and 95% confidence intervals, estimated using restricted maximum likelihood (REML) for each of our three datasets. Colors indicate the 11 variables included in each model (binary covariates for sampling season). Estimate confidence intervals are shaded by whether they cross zero (the vertical dashed line), with increased transparency denoting non-significant effects. The intercept contains the following reference levels: single sampling (sampling method); cross-sectional study (study format); single PCR (PCR type); fecal, rectal, or anal sample (tissue type); euthanasia not used (euthanasia use); Craseonycteridae (bat family); not fall, not winter, not spring, and not summer (sampling season); and RNA-dependent RNA polymerase (RdRp) only (gene target).

## Discussion

Since the onset of the COVID-19 pandemic, significantly increased research attention has been paid to bats as potential reservoir hosts of coronaviruses (including, presumably, many with zoonotic potential) [17–19]. While other studies have reported data on the geographical and taxonomic distribution of reported bat hosts [19,20], ours has generated the first standardized, PRISMA-generated open database of coronavirus surveillance in bats that provides disaggregated data (including negative results). In doing so, our study takes one of many first steps towards building an open database of wildlife disease surveillance with relevance to pandemic prediction and preparedness [21].

Our initial dataset represents a systematic snapshot of bat coronavirus research prior to the COVID-19 pandemic and includes 111 studies, 2,434 records, and a total of 93,877 bat samples. Our geographic and taxonomic analyses suggest a large focus on bat sampling in China compared to (and potentially at the expense of) gaps throughout South Asia, the Americas, Sub-Saharan Africa, and East Africa. Additionally, very few studies sampled in the United States and Canada (two and three, respectively). However, we acknowledge that progress towards addressing some of these gaps has been made since the onset of the pandemic; for example, more recent bat surveillance work has taken place in Latin America and Madagascar [19,22–26]. While phylogenetic coverage across bats is a strength of the dataset, we noted key taxonomic biases in the intensity of sampling efforts, with subclades of the Hipposideridae and Rhinolophidae families being some of the most heavily sampled taxa versus significant undersampling within the Stenodermatinae and Pteropodinae subfamilies. Priorities for future research should include strengthening surveillance efforts in these undersampled regions and bat taxa, especially as some have been predicted to harbor novel betacoronaviruses [19].

After controlling for bat phylogeny, sampling variance, and both study- and observation-level heterogeneity, repeat sampling and longitudinal study design were the only consistently significant predictors of positive coronavirus prevalence. Thus, to optimize detection sensitivity, substantial resources and careful planning should be allocated towards following this study format [27]. Additionally, euthanasia did not impact the likelihood of viral detection; thus, terminal sampling may not be necessary for studies attempting to detect coronavirus RNA, and our analysis suggests that coronavirus positivity will not be substantially biased by tissue or sample type. This is important for researchers, given that coronavirus surveillance can be accomplished with opportunistic (e.g., roost feces) and readily accessible (e.g., museum-derived) samples [28]. Further, avoiding euthanasia reduces negative impacts of virus surveillance studies on bat population dynamics, and also facilitates true longitudinal, mark-recapture designs.

Finally, our systematic data compilation process revealed marked challenges in synthesizing viral surveillance data from wildlife studies. Although study-level effects are in part accounted for with the random effects structure of our meta-analysis, we note that at least some of our non-significant results could still be due to variability in study format, sampling design, and reporting. To reduce this risk in future analyses, we encourage researchers collecting these data to be methodical in reporting their data at the finest resolution possible (i.e., fully stratified by location, timepoint, bat species, virus species or strain, tissue type, etc.). In the longer term, developing and adopting data standards for reporting these types of data—and developing real-time channels to aggregate them with standardized metadata—could significantly improve their ability to address key questions about transmission dynamics, bat immunology, viral evolution, and spillover risk.

## Methods

### Systematic review

To identify studies quantifying the proportion of wild bats positive for alpha- or betacoronaviruses using PCR or serological methods, we followed the Preferred Reporting Items for Systematic Reviews and Meta-Analyses (PRISMA) protocol (Figure S1) [29]. We systematically searched Web of Science, PubMed, and Global Health (a database comprising publications from the Public Health and Tropical Medicine database and CAB Abstracts). PubMed searches used the following string: (bat* OR Chiroptera*) AND (coronavirus* OR CoV*). Web of Science and Global Health (comprised of CAB Abstracts and Public Health and Tropical Medicine database) searches used the following string: (bat* OR Chiroptera*) AND (coronavirus* OR CoV*) AND (wild*). Searches were performed on September 24, 2020.

We screened a total of 1,016 abstracts for studies that included sampling of wild bats for coronaviruses. Publications were excluded if they did not assess coronavirus prevalence or seroprevalence in bats or were published in languages other than English. In total, we identified a total of 159 candidate articles that we screened for these data. Of these, 111 studies tested bats for coronaviruses, reported reusable data, and were included in our final, publicly available dataset. Geographic and taxonomic analyses, which did not rely on prevalence proportion positive, were performed on a 109-study subset of the public dataset which excludes records with genus- or family-level versus species-level bat data and includes seroprevalence data as well as data that could not be used to calculate prevalence (e.g., number of samples corresponds to geographic region rather than bat species). Infection prevalence analyses were performed on a 107-study subset of the public dataset. Each of these two datasets were then divided into three more: pooled-coronavirus genera, alphacoronavirus genus-only, and betacoronavirus genus-only (Table S1). The datasets used for geographic and taxonomic analyses, which included seroprevalence data as well as data that could not be used to calculate prevalence (e.g., number of samples corresponds to geographic region rather than bat species) had 176 (pooled-coronavirus genera), 56 (alphacoronavirus genus-only), and 143 (betacoronavirus genus-only) more rows than the corresponding infection prevalence datasets.

Our aim was to provide a comprehensive record of bat coronavirus surveillance up to the beginning of the COVID-19 pandemic, and our sample necessarily omits some more recent publications that have reanalyzed samples motivated by investigations into the evolutionary origins of SARS-CoV-2 and other L2 lineage sarbecoviruses. It also omits the final dataset compiled by the USAID PREDICT dataset and released at the end of 2020. While these data are an incomparable resource, their scope and standardized format makes them a substantively different kind of data than all other studies we analyze here; these data have been extensively analyzed elsewhere [1]. Perhaps most importantly, the majority of studies that report primary data on bat coronavirus testing by this program are included in our dataset.

### Data collection

Our initial dataset consists of a total of 111 studies and 2,434 records. Each record provides a prevalence or seroprevalence estimate at the finest spatiotemporal, methodological, and phylogenetic scale reported. More precisely, each unique record includes a distinct combination of coronavirus genus; bat genus, family, and/or species; sampled tissue; detection method (i.e., PCR or serology); gene/protein target; date, and geographic location (sampling country, state, and specific site and/or geographic coordinates, if available). Detection estimates derived at finer phylogenetic scales (e.g., virus strain) were aggregated to genus. As observed previously for bat filoviruses and henipaviruses, some studies pooled coronavirus detection estimates for more than one bat species [6]. Rows with these pooled prevalence estimates were excluded from subsequent statistical analyses. Sampling strategies were classified as longitudinal and cross-sectional: prevalence estimates derived from repeated sampling at one location were marked as longitudinal, while those derived from one location on a specific date were listed as cross-sectional. Thus, most studies (93.6%) yielded more than one detection estimate record: for example, a longitudinal study that provides individual coronavirus detection estimates from two types of tissue in a given bat species on six separate dates spanning several years would result in at least 12 records in the dataset.

In addition to these spatial and temporal components, we recorded data on detection methodology (e.g., single or nested/multiple PCR for RNA detection, ELISA for antibody detection, or immunohistochemistry), additional virus taxonomy (e.g., subgenus, strain), PCR primers (and their gene targets), and whether the authors included information on the sex of the sampled bats or the use of euthanasia.

### Geographic and taxonomic analyses of sampling effort

With these data, we assessed geographic and taxonomic patterns in bat sampling effort. For the former, we fit a generalized linear model (GLM) with whether a country had been sampled for bat coronaviruses as a binomial response and region as the predictor in R. For sampled countries (n=55), we fit equivalent GLMs that modeled the number of unique studies and the total samples per country as a Poisson-distributed response. For each GLM, we assessed fit using McFadden’s *R^2^* and the *performance* package [30]. We also adjusted for the inflated false-discovery rate in post-hoc comparisons using *emmeans* [31].

For taxonomic patterns, we derived equivalent response variables across bat species, using a recent phylogeny as a taxonomic backbone [13]. For all bat species in this phylogeny (*n* = 1287), we derived a binary response for whether a species had been sampled for coronaviruses. For those sampled species (*n* = 363), we derived the number of unique studies and the total samples. Using the *caper* package [32], we first estimated phylogenetic signal in sampling effort (i.e., the propensity for related bat species to be sampled in a similar intensity). For binary sampling effort, we calculated *D*, where a value of 1 indicates a phylogenetically random trait distribution and 0 indicates phylogenetic clustering under a Brownian motion model of evolution [33]. For sampled species, we estimated Pagel’s λ for the log_10_-transformed number of studies and samples [34]. Next, we applied a graph-partitioning algorithm, phylogenetic factorization, to more flexibly identify any bat clades across taxonomic levels that differ in sampling effort. With a standardized taxonomy from our bat phylogeny [13], we used the *phylofactor* package to partition binary sampling effort, number of studies, and number of samples in a series of iterative GLMs for each edge in the tree [14,35]. As in our geographic analyses, we modeled these variables with binomial and Poisson distributions. We then determined the number of significant clades using Holm’s sequentially rejective test with a 5% family-wise error rate [36].

### Phylogenetic meta-analysis of infection prevalence

We first used the *metafor* package to calculate Freeman–Tukey double arcsine transformed proportions of coronavirus infection-positive bats and their corresponding sampling variances [16 2010]. We then built two hierarchical meta-analysis models for three infection prevalence datasets: the global dataset, an alphacoronavirus-specific dataset, and a betacoronavirus-specific dataset (see Table S1 for the sample size per model). Each model was fit using restricted maximum likelihood and included bat species and phylogeny (using the previous bat tree) as random effects alongside an observation-level random effect nested within a study-level effect [15]. The first model (i.e., model 1) for each dataset only included an intercept and was used to estimate *I^2^*, which quantifies the contribution of true heterogeneity (rather than noise) to variance in infection prevalence [37]. We report both the overall *I^2^* per dataset as well as the proportional *I^2^* for each random effect, and we used Cochran’s *Q* to test if such heterogeneity was greater than that expected by sampling error alone. The second model (i.e., model 2) for each dataset included the following moderators: sampling method (repeat vs. single) study type (longitudinal vs. cross-sectional sampling), PCR type (nested/multiple vs. single), tissue analyzed, whether terminal sampling was performed, bat family, sampling season, and gene target. We calculated variance inflation factors of all moderators in the linear model: the moderators displayed no substantial collinearity [38]. To facilitate estimating model coefficients, we removed levels for any moderators with *n* < 3. For each iteration of model 2, we assessed moderator significance using the *Q* test (i.e., a Wald-like test of all coefficients per moderator) and estimated a pseudo-*R^2^* as the proportional reduction in the summed variance components compared against those from an intercept-only model [39].

## Acknowledgements

This work was supported by funding to the Viral Emergence Research Initiative (VERENA) consortium, including NSF BII 2021909, as well as by the National Institute of General Medical Sciences of the National Institutes of Health (P20GM134973).

## Author contributions

D.J.B., C.J.C., and L.E.C. devised the study. L.E.C., A.C.F., and B.C. performed the data collection. D.J.B. conducted the geographic and taxonomic analyses. L.E.C. conducted the phylogenetically controlled meta-analysis. L.E.C. and D.J.B. generated all figures and tables. L.E.C., A.C.F., C.J.C., and D.J.B. interpreted the results. L.E.C., A.C.F., C.J.C., and D.J.B. wrote the manuscript. All authors reviewed the manuscript and approved the submitted version.

## Data and code availability

The primary dataset is available on Github (www.github.com/viralemergence/datacov; DOI: 10.5281/zenodo.6644163). The unprocessed data and scripts to generate the primary dataset (and all other derived datasets) and to replicate all analyses and visualizations are available at www.github.com/viralemergence/batgap; DOI: 10.5281/zenodo.6644081).

## Figures and Tables

**Table 1.** Meta-analysis of coronavirus prevalence across studies. ANOVA table from the phylogenetic meta-analysis model fit using REML to all data and each data subset (alphacoronavirus only or betacoronavirus only). For each variable, we provide Cochran’s *Q*, the associated degrees of freedom, and the *p* value.

|  | any coronavirus genus |  |  | alphacoronavirus only |  |  | betacoronavirus only |  |  |
| --- | --- | --- | --- | --- | --- | --- | --- | --- | --- |
| | $Q$ | $df$ | $p$ | $Q$ | $df$ | $p$ | $Q$ | $df$ | $p$ |
| sampling method | 16.754 | 2 | < 0.001 | 9.516 | 2 | 0.009 | 18.765 | 2 | < 0.001 |
| study format | 6.650 | 1 | 0.01 | 7.283 | 1 | 0.007 | 2.380 | 1 | 0.123 |
| PCR type | 1.279 | 1 | 0.258 | 0.428 | 1 | 0.513 | 2.833 | 1 | 0.092 |
| tissue type | 36.536 | 8 | < 0.001 | 15.556 | 8 | 0.049 | 29.398 | 8 | < 0.001 |
| euthanasia use | 0.098 | 1 | 0.755 | 0.254 | 1 | 0.614 | 0.001 | 1 | 0.975 |
| bat family | 12.679 | 11 | 0.315 | 11.670 | 11 | 0.389 | 12.617 | 11 | 0.319 |
| sampling season | 8.406 | 4 | 0.078 | 10.177 | 4 | 0.038 | 7.263 | 11 | 0.123 |
| gene target | 1.989 | 2 | 0.370 | 0.556 | 2 | 0.758 | 2.408 | 2 | 0.300 |

## References

1. Anthony SJ, Johnson CK, Greig DJ, Kramer S, Che X, Wells H, et al. Global patterns in coronavirus diversity. Virus Evol. 2017;3.

2. Lednicky JA, Tagliamonte MS, White SK, Elbadry MA, Alam MM, Stephenson CJ, et al. Independent infections of porcine deltacoronavirus among Haitian children. Nature. 2021;600: 133–137.

3. Zhu Z, Lian X, Su X, Wu W, Marraro GA, Zeng Y. From SARS and MERS to COVID-19: a brief summary and comparison of severe acute respiratory infections caused by three highly pathogenic human coronaviruses. Respir Res. 2020;21: 1–14.

4. Woo PCY, Lau SKP, Li KSM, Poon RWS, Wong BHL, Tsoi H-W, et al. Molecular diversity of coronaviruses in bats. Virology. 2006;351: 180–187.

5. Woo PCY, Lau SKP, Lam CSF, Lau CCY, Tsang AKL, Lau JHN, et al. Discovery of seven novel Mammalian and avian coronaviruses in the genus deltacoronavirus supports bat coronaviruses as the gene source of alphacoronavirus and betacoronavirus and avian coronaviruses as the gene source of gammacoronavirus and deltacoronavirus. J Virol. 2012;86: 3995–4008.

6. Becker DJ, Crowley DE, Washburne AD, Plowright RK. Temporal and spatial limitations in global surveillance for bat filoviruses and henipaviruses. Biol Lett. 2019;15: 20190423.

7. Nusser SM, Clark WR, Otis DL, Huang L. Sampling considerations for disease surveillance in wildlife populations. Wildfire. 2008;72: 52–60.

8. Plowright RK, Eby P, Hudson PJ, Smith IL, Westcott D, Bryden WL, et al. Ecological dynamics of emerging bat virus spillover. Proc Biol Sci. 2015;282: 20142124.

9. Poon LLM, Chu DKW, Chan KH, Wong OK, Ellis TM, Leung YHC, et al. Identification of a novel coronavirus in bats. J Virol. 2005;79: 2001–2009.

10. Woo PCY, Lau SKP, Chu C-M, Chan K-H, Tsoi H-W, Huang Y, et al. Characterization and complete genome sequence of a novel coronavirus, coronavirus HKU1, from patients with pneumonia. J Virol. 2005;79: 884–895.

11. de Souza Luna LK, Heiser V, Regamey N, Panning M, Drexler JF, Mulangu S, et al. Generic detection of coronaviruses and differentiation at the prototype strain level by reverse transcription-PCR and nonfluorescent low-density microarray. J Clin Microbiol. 2007;45: 1049–1052.

12. Watanabe S, Masangkay JS, Nagata N, Morikawa S, Mizutani T, Fukushi S, et al. Bat coronaviruses and experimental infection of bats, the Philippines. Emerg Infect Dis. 2010;16: 1217–1223.

13. Upham NS, Esselstyn JA, Jetz W. Inferring the mammal tree: Species-level sets of phylogenies for questions in ecology, evolution, and conservation. PLoS Biol. 2019;17: e3000494.

14. Washburne AD, Silverman JD, Morton JT, Becker DJ, Crowley D, Mukherjee S, David LA, Plowright RK. Phylofactorization: a graph partitioning algorithm to identify phylogenetic scales of ecological data. Ecological Monographs. 2019 May;89(2):e01353.

15. Cinar O, Nakagawa S, Viechtbauer W. Phylogenetic multilevel meta-analysis: A simulation study on the importance of modelling the phylogeny. Methods Ecol Evol. 2022;13: 383–395.

16. Viechtbauer W. Conducting meta-analyses in R with the metafor package. Journal of Statistical Software. 2010;36(3):1–48.

17. Latinne A, Hu B, Olival KJ, Zhu G, Zhang L, Li H, et al. Origin and cross-species transmission of bat coronaviruses in China. Nat Commun. 2020;11: 4235.

18. Wacharapluesadee S, Tan CW, Maneeorn P, Duengkae P, Zhu F, Joyjinda Y, et al. Evidence for SARS-CoV-2 related coronaviruses circulating in bats and pangolins in Southeast Asia. Nat Commun. 2021;12: 972.

19. Becker DJ, Albery GF, Sjodin AR, Poisot T, Bergner LM, Chen B, et al. Optimising predictive models to prioritise viral discovery in zoonotic reservoirs. Lancet Microbe. 2022.

20. Ruiz-Aravena M, McKee C, Gamble A, Lunn T, Morris A, Snedden CE, et al. Ecology, evolution and spillover of coronaviruses from bats. Nat Rev Microbiol. 2022;20: 299–314.

21. The Verena Consortium. Building a global atlas of wildlife disease data. In: The Verena Blog. 2 Mar 2022. Available: https://www.viralemergence.org/blog/building-a-global-atlas-of-wildlife-disease-data

22. Alves RS, do Canto Olegário J, Weber MN, da Silva MS, Canova R, Sauthier JT, et al. Detection of coronavirus in vampire bats *(Desmodus rotundus)* in southern Brazil. Transbound Emerg Dis. 2021. doi:10.1111/tbed.14150

23. Bergner LM, Orton RJ, Streicker DG. Complete genome sequence of an alphacoronavirus from common vampire bats in Peru. Microbiol Resour Announc. 2020;9. doi:10.1128/MRA.00742-20

24. Becker DJ, Lei GS, Janech MG, Bland AM, Fenton MB, Simmons NB, Relich RF, Neely BA. Serum proteomics identifies immune pathways and candidate biomarkers of coronavirus infection in wild vampire bats. Frontiers in Virology. 2022; 2.

25. Kettenburg G, Kistler A, Ranaivoson HC, Ahyong V, Andrianiaina A, Andry S, et al. Full genome nobecovirus sequences from Malagasy fruit bats define a unique evolutionary history for this coronavirus clade. Front Public Health. 2022;10: 786060.

26. Hoarau AOG, Goodman SM, Al Halabi D, Ramasindrazana B, Lagadec E, Le Minter G, et al. Investigation of astrovirus, coronavirus and paramyxovirus co-infections in bats in the western Indian Ocean. Virol J. 2021;18: 205.

27. Plowright RK, Becker DJ, McCallum H, Manlove KR. Sampling to elucidate the dynamics of infections in reservoir hosts. Philos Trans R Soc Lond B Biol Sci. 2019;374: 20180336.

28. Thompson CW, Phelps KL, Allard MW, Cook JA, Dunnum JL, Ferguson AW, et al. Preserve a voucher specimen! The critical need for integrating natural history collections in infectious disease studies. MBio. 2021;12.

29. Moher D, Liberati A, Tetzlaff J, Altman DG. Preferred reporting items for systematic reviews and meta-analyses: the PRISMA statement. BMJ. 2009;339. doi:10.1136/bmj.b2535

30. Lüdecke D, Ben-Shachar M, Patil I, Waggoner P, Makowski D. *performance:* An R package for assessment, comparison and testing of statistical models. Journal of Open Source Software. 2021, 3139.

31. Benjamini Y, Hochberg Y. Controlling the false discovery rate: A practical and powerful approach to multiple testing. J R Stat Soc. 1995;57: 289–300.

32. Orme D, Freckleton R, Thomas G, Petzoldt T, Fritz S, Isaac N, et al. caper: comparative analyses of phylogenetics and evolution in R. 2012;2: 458.

33. Fritz SA, Purvis A. Phylogenetic diversity does not capture body size variation at risk in the world’s mammals. Proc Biol Sci. 2010;277: 2435–2441.

34. Pagel M. Inferring the historical patterns of biological evolution. Nature. 1999;401: 877–884.

35. Crowley D, Becker D, Washburne A, Plowright R. Identifying suspect bat reservoirs of emerging infections. Vaccines. 2020; 8.

36. Holm S. A simple sequentially rejective multiple test procedure. Scand Stat Theory Appl. 1979;6: 65–70.

37. Senior AM, Grueber CE, Kamiya T, Lagisz M, O’Dwyer K, Santos ESA, et al. Heterogeneity in ecological and evolutionary meta-analyses: its magnitude and implications. Ecology. 2016;97: 3293–3299.

38. Zuur AF, Ieno EN, Elphick CS. A protocol for data exploration to avoid common statistical problems. Methods Ecol Evol. 2010;1: 3–14.

39. López-López JA, Marín-Martínez F, Sánchez-Meca J, Van den Noortgate W, Viechtbauer W. Estimation of the predictive power of the model in mixed-effects meta-regression: A simulation study. Br J Math Stat Psychol. 2014;67: 30–48.

