## Supporting information for "Sampling strategies and pre-pandemic surveillance gaps for bat coronaviruses"

**Figure S1.** PRISMA reporting for systematic review and meta-analysis.

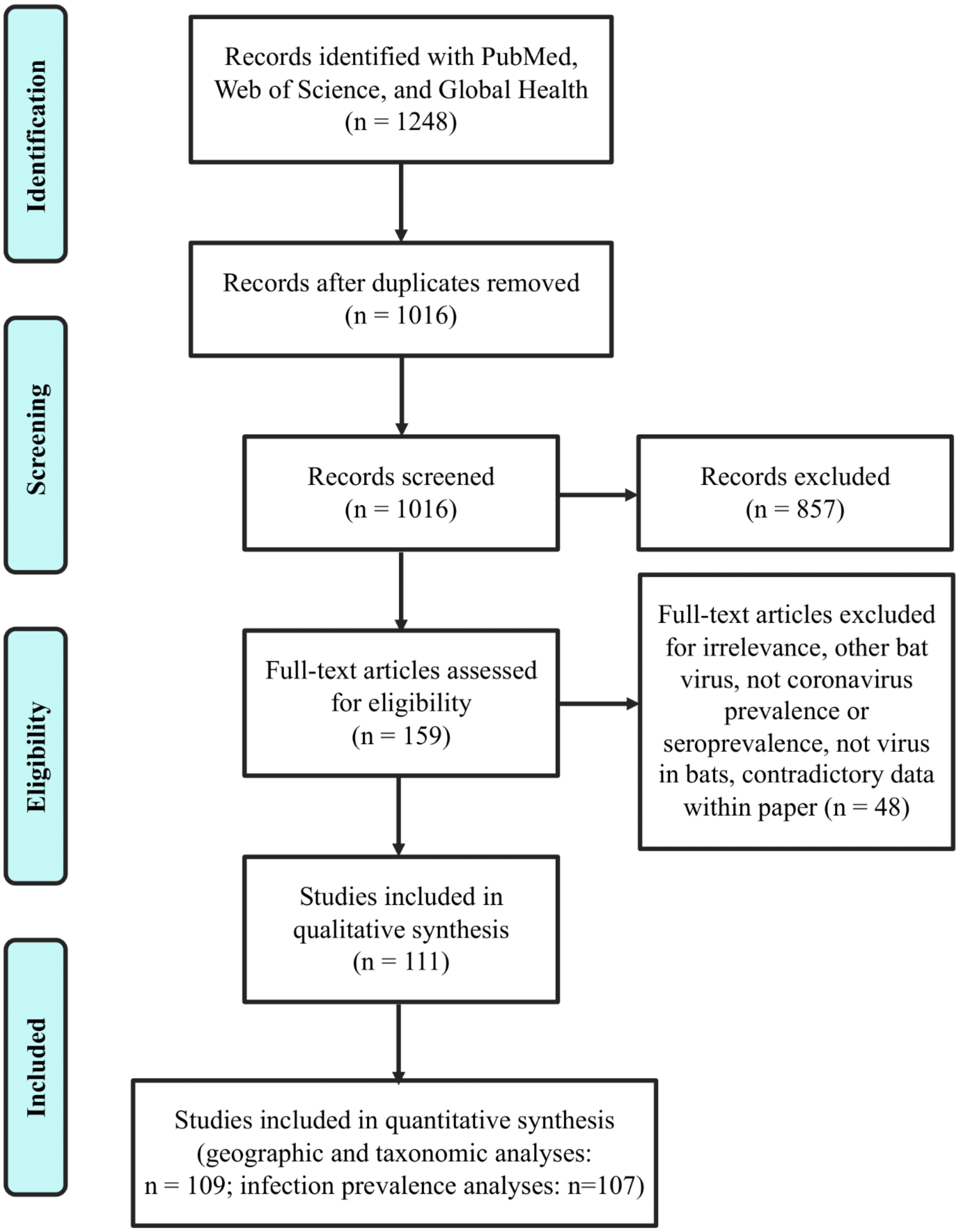

**Table S1.** Data included in each base meta-analytic model (fit using REML, intercept only). Levels of factors which appeared fewer than three times in the datasets (i.e., liver [tissue type]; Noctilionidae and Mystacinidae [bat family]) were not included in the full meta-analysis model with reported coefficients (fit using REML, see Figure 3).

| **Viral genus (as reported by study)** | **Rows of available data** | | **Included in models:** | | |
|---|---|---|---|---|---|
| | **# rows (non-zero)** | **# rows (zero only)** | **CoV** | **α** | **β** |
| alphacoronavirus prevalence (explicit) | 407 | 20 | ✓ | ✓ | x |
| betacoronavirus prevalence (explicit) | 315 | 56 | ✓ | x | ✓ |
| genus unspecified or unresolved<br><br>(incl. if tests noted to include alphas and betas, but not specific, or if general sampling noted to include alpha and/or beta but not reported as proportion of sample) | 56 | 1400* | ✓ | 0 | 0 |
| coinfection of any kind | 15 | 0 | x | x | x |

*These rows comprise the zero prevalence estimates from studies using pan-coronavirus primers (most studies).

**Table S2.** Distribution of tissue types and associated percentages of positive and zero infection prevalence values.

| | any coronavirus genus | | | alphacoronavirus only | | | betacoronavirus only | | |
|---|---|---|---|---|---|---|---|---|---|
| tissue type | *n* | % positive prevalence | % zero prevalence | n | % positive prevalence | % zero prevalence | *n* | % positive prevalence | % zero prevalence |
| fecal, rectal, or anal | 1084 | 39.21 | 60.79 | 895 | 28.16 | 71.84 | 791 | 19.09 | 80.91 |
| liver | 2 | 100 | 0 | 0 | - | - | 2 | 100 | 0 |
| blood or serum | 3 | 0 | 100 | 3 | 0 | 100 | 3 | 0 | 100 |
| intestine | 183 | 39.34 | 60.66 | 144 | 22.92 | 77.08 | 144 | 22.92 | 77.08 |
| lung or respiratory | 74 | 9.46 | 90.54 | 66 | 6.06 | 93.94 | 70 | 4.29 | 95.71 |
| oropharyngeal | 131 | 6.11 | 93.89 | 126 | 5.56 | 94.44 | 123 | 0.81 | 99.19 |
| skin swab | 11 | 0 | 100 | 11 | 0 | 100 | 11 | 0 | 100 |
| urinary | 55 | 3.64 | 96.36 | 55 | 3.64 | 96.36 | 53 | 0 | 100 |
| pooled swabs/samples | 284 | 37.32 | 62.68 | 234 | 24.36 | 75.64 | 212 | 16.04 | 83.96 |
| pooled tissue | 58 | 24.14 | 75.86 | 53 | 16.98 | 83.02 | 49 | 10.20 | 89.80 |

**Table S3.** Post-hoc GLM results (binomial errors) for binary sampling effort across countries globally as a function of geography. The *p*-values are adjusted with the Benjamini-Hochberg correction.

| | Comparison | Odds ratio | SE | z ratio | *p* |
|---|---|---|---|---|---|
| 1 | Africa / Americas | 2.83 | 1.49 | 1.99 | 0.12 |
| 2 | Africa / Asia | 0.73 | 0.31 | -0.74 | 0.65 |
| 3 | Africa / Europe | 0.8 | 0.34 | -0.52 | 0.75 |
| 4 | Africa / Oceania | 2.89 | 1.96 | 1.56 | 0.20 |
| 5 | Americas / Asia | 0.26 | 0.14 | -2.58 | 0.08 |
| 6 | Americas / Europe | 0.28 | 0.15 | -2.39 | 0.08 |
| 7 | Americas / Oceania | 1.02 | 0.76 | 0.03 | 0.98 |
| 8 | Asia / Europe | 1.1 | 0.47 | 0.22 | 0.92 |
| 9 | Asia / Oceania | 3.96 | 2.7 | 2.02 | 0.12 |
| 10 | Europe / Oceania | 3.61 | 2.47 | 1.88 | 0.12 |

**Table S4.** Post-hoc GLM results (Poisson errors) for the number of studies per country as a function of geography across all sampled countries. The *p*-values are adjusted with the Benjamini-Hochberg correction.

| | **Comparison** | **Risk ratio** | **SE** | **z ratio** | ***p*** |
|---|---|---|---|---|---|
| 1 | Africa / Americas | 0.71 | 0.25 | -1 | 0.64 |
| 2 | Africa / Asia | 0.42 | 0.1 | -3.57 | < 0.001 |
| 3 | Africa / Europe | 0.82 | 0.23 | -0.7 | 0.80 |
| 4 | Africa / Oceania | 0.77 | 0.35 | -0.58 | 0.80 |
| 5 | Americas / Asia | 0.59 | 0.18 | -1.74 | 0.28 |
| 6 | Americas / Europe | 1.16 | 0.39 | 0.44 | 0.82 |
| 7 | Americas / Oceania | 1.08 | 0.53 | 0.16 | 0.88 |
| 8 | Asia / Europe | 1.98 | 0.45 | 2.97 | 0.02 |
| 9 | Asia / Oceania | 1.84 | 0.79 | 1.43 | 0.38 |
| 10 | Europe / Oceania | 0.93 | 0.42 | -0.15 | 0.88 |

**Table S5.** Post-hoc GLM results (Poisson errors) for the number of samples tested per country as a function of geography across all sampled countries. The *p*-values are adjusted with the Benjamini-Hochberg correction.

| | **Comparison** | **Risk ratio** | **SE** | **z ratio** | ***p*** |
|---|---|---|---|---|---|
| 1 | Africa / Americas | 0.89 | 0.01 | -7.42 | < 0.001 |
| 2 | Africa / Asia | 0.43 | 0 | -84.85 | < 0.001 |
| 3 | Africa / Europe | 1.88 | 0.03 | 44 | < 0.001 |
| 4 | Africa / Oceania | 0.62 | 0.01 | -27.54 | < 0.001 |
| 5 | Americas / Asia | 0.48 | 0.01 | -53.66 | < 0.001 |
| 6 | Americas / Europe | 2.11 | 0.04 | 43.4 | < 0.001 |
| 7 | Americas / Oceania | 0.7 | 0.01 | -18.39 | < 0.001 |
| 8 | Asia / Europe | 4.41 | 0.06 | 115.9 | < 0.001 |
| 9 | Asia / Oceania | 1.46 | 0.02 | 23.76 | < 0.001 |
| 10 | Europe / Oceania | 0.33 | 0.01 | -58.41 | < 0.001 |

**Table S6.** Number of studies, number of samples analyzed, and percentage of dataset by global region.

| | any coronavirus genus | | | alphacoronavirus only | | | betacoronavirus only | | |
|---|---|---|---|---|---|---|---|---|---|
| global region | number of studies | number of samples analyzed | percentage of dataset (%) | number of studies | number of samples analyzed | percentage of dataset (%) | number of studies | number of samples analyzed | percentage of dataset (%) |
| Africa | 17 | 357 | 17.2 | 14 | 275 | 15.5 | 17 | 286 | 17.6 |
| Americas | 12 | 352 | 17.0 | 11 | 341 | 19.3 | 8 | 276 | 17.0 |
| Asia | 54 | 876 | 42.2 | 43 | 712 | 40.2 | 51 | 710 | 43.6 |
| Australia or New Zealand | 5 | 36 | 1.7 | 5 | 34 | 1.9 | 2 | 18 | 1.1 |
| Europe | 24 | 455 | 21.9 | 23 | 408 | 23.1 | 23 | 337 | 20.7 |

**Table S7.** Number of studies, number of samples analyzed, and percentage of dataset by global region and country.

| | any coronavirus genus | | | alphacoronavirus only | | | betacoronavirus only | | |
|---|---|---|---|---|---|---|---|---|---|
| global region: country | number of studies | number of samples analyzed | percentage of dataset (%) | number of studies | number of samples analyzed | percentage of dataset (%) | number of studies | number of samples analyzed | percentage of dataset (%) |
| Africa: Egypt | 2 | 11 | 0.5 | 1 | 1 | 0.1 | 2 | 11 | 0.7 |
| Africa: Gabon | 1 | 40 | 1.9 | 1 | 37 | 2.1 | 1 | 34 | 2.1 |
| Africa: Ghana | 3 | 39 | 1.9 | 3 | 34 | 1.9 | 3 | 35 | 2.2 |
| Africa: Guinea | 1 | 45 | 2.2 | 1 | 28 | 1.6 | 1 | 36 | 2.2 |
| Africa: Kenya | 3 | 81 | 3.9 | 3 | 57 | 3.2 | 3 | 48 | 3.0 |
| Africa: Madagascar | 2 | 45 | 2.2 | 2 | 37 | 2.1 | 2 | 41 | 2.5 |
| Africa: Mauritius | 1 | 3 | 0.2 | 1 | 3 | 0.2 | 1 | 3 | 0.2 |
| Africa: Morocco | 1 | 4 | 0.2 | 1 | 4 | 0.2 | 1 | 3 | 0.2 |
| Africa: Mozambique | 1 | 10 | 0.5 | 1 | 9 | 0.5 | 1 | 4 | 0.3 |
| Africa: Nigeria | 1 | 1 | 0.1 | - | - | - | 1 | 1 | 0.1 |
| Africa: Rwanda | 1 | 49 | 2.4 | 1 | 37 | 2.1 | 1 | 46 | 2.8 |
| Africa: Seychelles | 1 | 1 | 0.1 | 1 | 1 | 0.1 | 1 | 1 | 0.1 |
| Africa: South Africa | 2 | 25 | 1.2 | 1 | 24 | 1.4 | 2 | 22 | 1.4 |
| Africa: Tunisia | 1 | 3 | 0.2 | 1 | 3 | 0.2 | 1 | 1 | 0.1 |
| Americas: Brazil | 4 | 101 | 4.9 | 3 | 97 | 5.5 | 3 | 74 | 4.5 |
| Americas: Canada | 3 | 6 | 0.3 | 3 | 6 | 0.3 | - | - | - |
| Americas: Costa Rica | 1 | 41 | 2.0 | 1 | 41 | 2.3 | 1 | 37 | 2.3 |
| Americas: Mexico | 1 | 72 | 3.5 | 1 | 75 | 4.2 | 1 | 71 | 4.4 |
| Americas: Trinidad | 1 | 14 | 0.7 | 1 | 14 | 0.8 | 1 | 12 | 0.7 |

| | | | | | | | | | |
|---|---|---|---|---|---|---|---|---|---|
| Americas: USA | 2 | 108 | 5.2 | 2 | 108 | 6.1 | 2 | 82 | 5.0 |
| Asia: China | 32 | 491 | 23.7 | 22 | 378 | 21.4 | 31 | 375 | 23.1 |
| Asia: Indonesia | 2 | 7 | 0.3 | 2 | 6 | 0.3 | 2 | 7 | 0.4 |
| Asia: Japan | 2 | 9 | 0.4 | 2 | 9 | 0.5 | 1 | 3 | 0.2 |
| Asia: Korea | 1 | 5 | 0.2 | 1 | 4 | 0.2 | 1 | 4 | 0.3 |
| Asia: Lebanon | 1 | 4 | 0.2 | 1 | 2 | 0.1 | 1 | 4 | 0.3 |
| Asia: Malaysian Borneo | 1 | 23 | 1.1 | 1 | 20 | 1.1 | 1 | 20 | 1.2 |
| Asia: Myanmar | 1 | 12 | 0.6 | 1 | 12 | 0.7 | 1 | 11 | 0.7 |
| Asia: Philippines | 2 | 43 | 2.1 | 2 | 21 | 1.2 | 2 | 41 | 2.5 |
| Asia: Saudi Arabia | 1 | 33 | 1.6 | 1 | 30 | 1.7 | 1 | 28 | 1.7 |
| Asia: Singapore | 1 | 6 | 0.3 | 1 | 6 | 0.3 | 1 | 6 | 0.4 |
| Asia: South Korea | 1 | 66 | 3.2 | 1 | 66 | 3.7 | 1 | 66 | 4.1 |
| Asia: Sri Lanka | 1 | 1 | 0.1 | - | - | - | 1 | 1 | 0.1 |
| Asia: Taiwan | 4 | 29 | 1.4 | 4 | 19 | 1.1 | 3 | 12 | 0.7 |
| Asia: Thailand | 3 | 143 | 6.9 | 3 | 135 | 7.6 | 3 | 129 | 7.9 |
| Asia: Vietnam | 2 | 4 | 0.2 | 2 | 4 | 0.2 | 2 | 3 | 0.2 |
| Australia | 4 | 35 | 1.7 | 4 | 33 | 1.9 | 2 | 18 | 1.1 |
| New Zealand | 1 | 1 | 0.1 | 1 | 1 | 0.1 | - | - | - |
| Europe: Belgium | 1 | 9 | 0.4 | 1 | 9 | 0.5 | 1 | 9 | 0.6 |
| Europe: Bulgaria | 1 | 23 | 1.1 | 1 | 19 | 1.1 | 1 | 17 | 1.1 |
| Europe: Denmark | 1 | 42 | 2.0 | 1 | 41 | 2.3 | 1 | 26 | 1.6 |
| Europe: Finland | 1 | 8 | 0.4 | 1 | 7 | 0.4 | 1 | 5 | 0.3 |
| Europe: France | 4 | 113 | 5.4 | 4 | 110 | 6.2 | 4 | 95 | 5.8 |
| Europe: Germany | 4 | 32 | 1.5 | 4 | 32 | 1.8 | 3 | 11 | 0.7 |

| | | | | | | | | | |
|---|---|---|---|---|---|---|---|---|---|
| Europe: Hungary | 1 | 24 | 1.2 | 1 | 23 | 1.3 | 1 | 18 | 1.1 |
| Europe: Italy | 6 | 89 | 4.3 | 5 | 64 | 3.6 | 6 | 69 | 4.2 |
| Europe: Luxembourg | 1 | 6 | 0.3 | 1 | 4 | 0.2 | 1 | 4 | 0.3 |
| Europe: Netherlands | 2 | 15 | 0.7 | 2 | 14 | 0.8 | 2 | 11 | 0.7 |
| Europe: Romania | 1 | 3 | 0.2 | - | - | - | 1 | 3 | 0.2 |
| Europe: Slovenia | 1 | 7 | 0.4 | 1 | 6 | 0.3 | 1 | 7 | 0.4 |
| Europe: Spain | 2 | 73 | 3.5 | 2 | 69 | 3.8 | 2 | 54 | 3.3 |
| Europe: Ukraine | 1 | 3 | 0.2 | 1 | 2 | 0.1 | 1 | 3 | 0.2 |
| Europe: UK | 1 | 8 | 0.4 | 1 | 8 | 0.5 | 1 | 5 | 0.2 |

**Table S8.** Results of phylogenetic factorization applied to the number of studies per sampled bat species (Poisson GLM). The number of retained phylogenetic factors (following a 5% family-wise error rate applied to GLMs), taxa corresponding to those clades, number of species, and mean number of studies for the clade compared to the paraphyletic remainder are shown. Taxonomy follows Upham et al. 2019.

| | **factor** | **taxa** | **tips** | **node** | **clade** | **other** |
|---|---|---|---|---|---|---|
| 1 | 1 | *Myotis escalerai, Myotis pequinius, Myotis chinensis, Myotis nattereri, Myotis punicus, Myotis myotis, Myotis blythii, Myotis bechsteinii, Myotis petax, Myotis macrodactylus, Myotis fimbriatus, Myotis pilosus, Myotis daubentonii* | 13 | 522 | 6.9 | 2.8 |
| 2 | 2 | *Pipistrellus nanulus, Pipistrellus nathusii, Pipistrellus pygmaeus, Pipistrellus pipistrellus, Pipistrellus hesperidus, Pipistrellus kuhlii, Pipistrellus deserti, Pipistrellus tenuis, Pipistrellus javanicus, Pipistrellus abramus, Nyctalus* | 15 | 415 | 6.0 | 2.8 |
| 3 | 3 | Rhinolophidae, Hipposideridae | 62 | 619 | 3.9 | 2.7 |
| 4 | 4 | Miniopterus | 16 | 546 | 4.1 | 2.9 |

**Table S9.** Results of phylogenetic factorization applied to the number of tested samples per sampled bat species (Poisson GLM). The number of retained phylogenetic factors (following a 5% family-wise error rate applied to GLMs), taxa corresponding to those clades, number of species, and mean number of tested samples for the clade compared to the paraphyletic remainder are shown. Taxonomy follows Upham et al. 2019.

| | factor | taxa | tips | node | clade | other |
|---|---|---|---|---|---|---|
| 1 | 1 | *Rhinolophus osgoodi, Rhinolophus affinis, Rhinolophus coelophyllus, Rhinolophus shameli, Rhinolophus stheno, Rhinolophus rouxii, Rhinolophus thomasi, Rhinolophus sinicus, Rhinolophus megaphyllus, Rhinolophus borneensis, Rhinolophus lepidus, Rhinolophus pusillus, Rhinolophus rex, Rhinolophus macrotis, Rhinolophus malayanus, Rhinolophus rufus* | 16 | 637 | 1568.1 | 182.3 |
| 2 | 2 | *Hipposideros cervinus, Hipposideros pomona, Hipposideros cineraceus, Hipposideros ater, Hipposideros halophyllus, Hipposideros dyacorum, Hipposideros ruber, Hipposideros abae, Hipposideros caffer, Hipposideros fuliginosus* | 10 | 663 | 979.4 | 222.5 |
| 3 | 3 | Miniopterus | 16 | 546 | 520.3 | 230.6 |
| 4 | 4 | Eonycteris, Rousettus, Epomops, Epomophorus, Micropteropus, Nanonycteris, Hypsignathus, Myonycteris | 16 | 705 | 503.5 | 231.4 |
| 5 | 5 | *Myotis escalerai, Myotis pequinius, Myotis chinensis, Myotis nattereri, Myotis punicus, Myotis myotis, Myotis blythii, Myotis bechsteinii, Myotis petax, Myotis macrodactylus, Myotis fimbriatus, Myotis pilosus, Myotis daubentonii* | 13 | 522 | 398.8 | 237.6 |
| 6 | 6 | Rhinolophidae, Hipposideridae | 36 | 619 | 217.7 | 246.2 |
| 7 | 7 | *Pipistrellus nanulus, Pipistrellus nathusii, Pipistrellus pygmaeus, Pipistrellus pipistrellus, Pipistrellus hesperidus, Pipistrellus kuhlii, Pipistrellus deserti, Pipistrellus tenuis, Pipistrellus javanicus, Pipistrellus abramus,* Nyctalus | 15 | 415 | 234.5 | 243.8 |
| 8 | 8 | Centurio, Artibeus, Dermanura, Vampyressa, Platyrrhinus, Uroderma, Sturnira, Carollia | 21 | 579 | 175.1 | 247.6 |
| 9 | 9 | Eidolon, Pteropus, Acerodon | 17 | 688 | 167.8 | 247.1 |

| | | | | | | |
|---|---|---|---|---|---|---|
| 10 | 10 | Glossophaga, Leptonycteris, Anoura, Hylonycteris, Choeroniscus, Trachops, Mimon, Phyllostomus, Tonatia, Phylloderma, Lophostoma, Chrotopterus | 16 | 569 | 12.2 | 254.0 |
| 11 | 11 | Corynorhinus, Euderma, Plecotus, Barbastella, Rhogeessa, Bauerus, Antrozous, Lasiurus, Parastrellus | 18 | 465 | 25.9 | 254.7 |
| 12 | 12 | *Rhinolophus capensis, Rhinolophus simulator, Rhinolophus denti, Rhinolophus blasii, Rhinolophus euryale, Rhinolophus mehelyi, Rhinolophus mossambicus, Rhinolophus fumigatus, Rhinolophus hildebrandtii, Rhinolophus darlingi, Rhinolophus ferrumequinum, Rhinolophus clivosus* | 12 | 623 | 318.0 | 240.8 |
| 13 | 13 | Coelops, Aselliscus, Hipposideros | 12 | 659 | 247.3 | 243.3 |
| 14 | 14 | Scoteanax, Kerivoula, Harpiola, Murina, Scotorepens | 12 | 488 | 21.1 | 251.0 |
| 15 | 15 | Neoromicia, Pipistrellus, Falsistrellus, Nyctophilus | 14 | 434 | 34.1 | 251.8 |
| 16 | 16 | Tylonycteris, Nycticeinops, Vespadelus, Chalinolobus | 10 | 431 | 198.8 | 244.7 |
| 17 | 17 | Scotophilus, Vespertilio, Glauconycteris, Eptesicus, Lasionycteris, Ia, Scotomanes | 21 | 410 | 111.5 | 251.5 |
| 18 | 18 | *Myotis aurascens, Myotis ikonnikovi, Myotis altarium, Myotis dasycneme, Myotis muricola, Myotis secundus, Myotis occultus, Myotis horsfieldii, Myotis macropus, Myotis adversus, Myotis capaccinii, Myotis longipes, Myotis laniger, Myotis siligorensis, Myotis davidii* | 15 | 519 | 48.9 | 251.8 |
| 19 | 19 | Mystacinidae, Noctilionidae, Mormoopidae, Phyllostomidae | 13 | 561 | 47.8 | 250.7 |
| 20 | 20 | *Mormopterus beccarii, Mormopterus norfolkensis, Tadarida brasiliensis, Otomops, Nyctinomops, Molossops, Cynomops, Molossus, Eumops, Tadarida teniotis, Tadarida australis, Mormopterus acetabulosus* | 17 | 391 | 55.4 | 252.6 |
| 21 | 21 | *Myotis mystacinus, Myotis alcathoe, Myotis elegans, Myotis riparius, Myotis keaysi, Myotis yumanensis, Myotis velifer, Myotis nigricans, Myotis brandtii, Myotis volans, Myotis ciliolabrum, Myotis californicus, Myotis lucifugus, Myotis evotis, Myotis thysanodes* | 15 | 501 | 63.6 | 251.1 |
| 22 | 22 | Pteropodidae | 14 | 680 | 77.5 | 250.0 |
| 23 | 23 | Molossidae | 10 | 382 | 119.4 | 246.9 |
